# HiChIP-Peaks: A HiChIP peak calling algorithm

**DOI:** 10.1101/682781

**Authors:** Chenfu Shi, Magnus Rattray, Gisela Orozco

## Abstract

**Motivation:** HiChIP is a powerful tool to interrogate 3D chromatin organization. Current tools to analyse chromatin looping mechanisms using HiChIP data require the identification of loop anchors to work properly. However, current approaches to discover these anchors from HiChIP data are not satisfactory, having either a very high false discovery rate or strong dependence on sequencing depth. Moreover, these tools do not allow quantitative comparison of peaks across different samples, failing to fully exploit the information available from HiChIP datasets.

**Results:** We develop a new tool based on a representation of HiChIP data centred on the re-ligation sites to identify peaks from HiChIP datasets, which can subsequently be used in other tools for loop discovery. This increases the reliability of these tools and improves recall rate as sequencing depth is reduced. We also provide a method to count reads mapping to peaks across samples, which can be used for differential peak analysis using HiChIP data.

**Availability:** HiChIP-Peaks is freely available at https://github.com/ChenfuShi/HiChIP_peaks

## 1 Introduction

The three-dimensional conformation of the chromatin is fundamental in the regulation of gene expression; regulatory elements such as enhancers have been shown to act by physically interacting with their target promoters (Yao *et al*., 2015; Shlyueva *et al*., 2014; Bulger and Groudine, 2011; Nolis *et al*., 2009). These regulatory elements are highly regulated and context specific (Simeonov *et al*., 2017; Kundaje *et al*., 2015; Alasoo *et al*., 2018). However, the requirement of large number of cells (tens of millions) to obtain chromatin interactions maps at sufficient resolution and the high cost associated with widely-used chromatin conformation techniques such as Hi-C have hindered the study of chromatin interactions in primary and patient-derived cells (Rao *et al*., 2014).

HiChIP is a recently developed technique to analyse chromatin conformation which consists of an in-situ Hi-C library preparation followed by a chromatin immunoprecipitation (ChIP) step, usually targeting the histone modification H3K27ac or cohesin. It has many advantages compared to traditional methods such as Hi-C, ChIA-PET and Capture Hi-C such as lower cost, higher sensitivity, lower input requirements and reduced sequencing required (Mumbach *et al*., 2016, 2017). Unfortunately, few tools exist to specifically analyse HiChIP data, with most publications relying on tools originally developed for Hi-C. HiChIP provides a new set of computational challenges because it combines biases introduced by two independent techniques: ChIP and in-situ Hi-C library preparation. This phenomenon is particularly evidenced by libraries enriched for H3K27ac because this histone modification has a significantly more specific enrichment compared to cohesin.

It is theoretically possible to extract two types of information from HiChIP data: the position of enriched regions for the chromatin immunoprecipitation and the long-range interactions involving these regions. The enriched regions, also called anchors or peaks, are usually identified prior to the identification of long-range interactions. Previous tools used either MACS2 on close range read pairs (fitHiChIP) (Bhattacharyya *et al*., 2018) or an adaptation of it (hichipper) (Lareau and Aryee, 2018). Vanilla MACS2 (Zhang *et al*., 2008) and its implementation in fitHiChIP has been shown to be strongly biased due to HiChIP specific biases, primarily the biotin pulldown (Lareau and Aryee, 2018) (Fig. S1). Hichipper tries to solve this problem by modelling a corrected background as a function of proximity to restriction sites and using that background for MACS2 peak calling. This results in many small peaks which then need to be merged to match the restriction fragments, which causes them to lose statistical metrics such as p values or scores rendering comparisons between samples infeasible. Our tests show also that using all the reads results in poor specificity while using only self-circle and dangling end reads results in very few reads being retained and correspondingly reduced sensitivity.

For this reason, many recent publications used independent ChIP-Seq as input to define anchors (Pelikan *et al*., 2018). However, that too can be a problem because the peaks definition can strongly influence the expected signal from a region and can be extremely variable if not done from exactly the same sample.

Here we propose a method to extract the location of ChIP-Seq peaks from HiChIP data that improves significantly on previous attempts. We analysed the HiChIP protocol and library preparation and developed an algorithm and a data representation that takes in consideration how the libraries are generated and Hi-C and HiChIP specific biases such as the biotin pulldown. We opted for a re-ligation (restriction) site centred representation and we model the expected background signal as an over dispersed Poisson/negative binomial model. We show that our approach is highly reproducible when compared to reference ChIP-Seq datasets and we show how this can improve the performance of downstream tools to call chromatin loops from HiChIP data. We also provide a method to count reads mapping to peaks across samples, which can be used to analyse differentially bound regions from HiChIP data.

The software is available as a Python 3 package on GitHub and PyPi along with code to reproduce the results presented here.

## 2 Methods

### 2.1 A novel representation for HiChIP data

Hi-C maps have typically been analysed using a fixed size bin matrix format. This can introduce significant biases because the expected number of reads depends on the number of restriction sites included within each bin. Moreover, reads are not uniformly distributed in the genome but are strongly biased around restriction sites because of the on-bead library preparation (Fig. S1). This causes sparsity and non-uniformity in the data, which can bias methods based on genomic position alone. An alternative approach is to analyse maps at a restriction fragment resolution. Analysing the raw data from Mumbach et al. (2017) we find that a significant number of reads contain uncut restriction sites (Table S1). This suggests that the cutting frequency is low and that especially for libraries prepared using frequently cutting enzymes (4-cutters) such as MboI the read assignment to the restriction fragment can be misleading. Moreover, traditional pair classifications such as dangling end and self-circle can be misleading since they can be wrongly classified as valid pairs (Fig. 1B) with significant biases due to fragment size (Fig. 1C). Importantly, the low frequency of cutting implies that the detectable signal is directly correlated to the number of restriction sites and only indirectly to the effective genomic size.

**Fig. 1.**
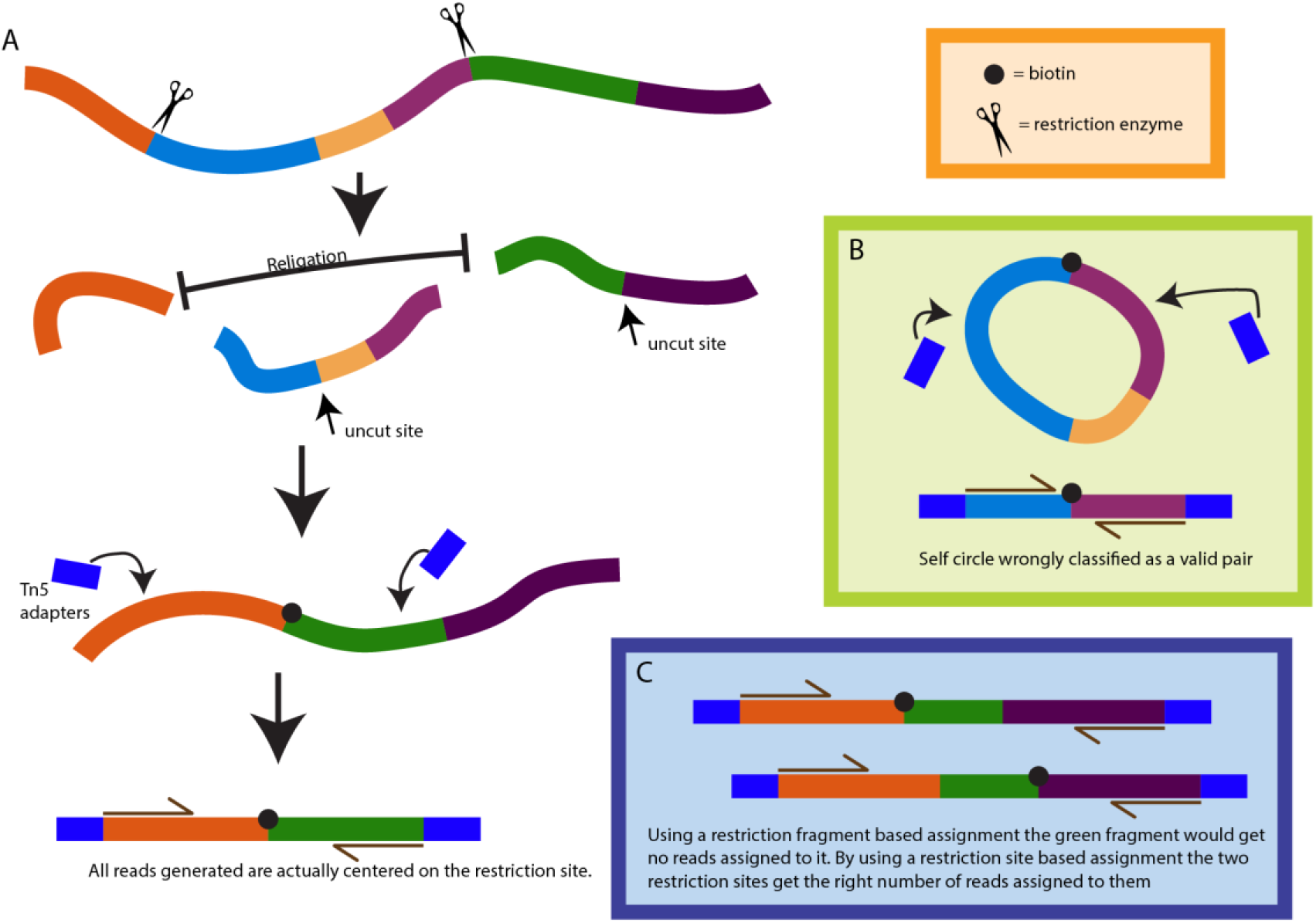
Justification for re-ligation site based data structure. A) The Hi-C protocol creates reads that are centered on the re-ligation site. Mapping reads at a higher resolution is not biologically relevant and only creates sparsity. B) Example of how traditional self-circle classifications are not reliable with libraries generated using frequently cutting enzymes. C) Example of how a data representation based on the restriction fragment can heavily bias the read counts depending on the size of the fragment while basing the data representation on the re-ligation site can reduce the bias by compensating each read assigned incorrectly with another one.

Approaching the problem in a novel way, we develop a data structure that focuses on the re-ligation site. This is the location from which reads are generated during the library preparation (Fig. 1A) and this data structure maximizes our detection power while at the same time minimizing biases introduced by the Hi-C library preparation. Reads are assigned based on the direction of the read to the nearest restriction site to which they point. This method significantly reduces the previously mentioned biases and maximizes information at the highest meaningful resolution. Miss-assignment of the reads due to small fragment size would be automatically corrected in a logical way (Fig. 1C). We implement this data structure as a sparse matrix in which the diagonal contains all the re-ligation pairs and the diagonal +1 contains pairs traditionally classified as self-circle and dangling ends. Starting from this data structure we develop our peak calling algorithm.

### 2.2 Peak calling

To limit the bias in the definition of self-circles and dangling ends and increase sensitivity we decided to also include reads that map to close range interactions. By default, we include interactions that are within two sites of the re-ligation pairs. We perform a moving integration as a smoothing function over three restriction sites to reduce noise. This allows us to use more significant reads in the successive steps and to regulate these settings depending on the input data. For example, libraries generated using the commercially available Arima-HiC kit, which uses two restriction enzymes, generate next to no reported self-circle and dangling ends reads (Fig. S2) but can be easily tuned by changing the previous parameters.

Visual inspection of the distribution of the background signal suggests that it closely matches a negative binomial distribution (Fig. S3), similar to what has been found with ChIP-seq data (Diaz *et al*., 2012). In order to model the distribution of the background it is important to note that a large majority of reads will locate in peaks (up to 80%) and inclusion of these reads would highly bias parameter estimation. We therefore first remove the most significant peaks using a Poisson based model (similarly to MACS2) (Zhang *et al*., 2008) with a very stringent setting (p-value < 1×10^−8^). We then estimate the negative binomial mean and over-dispersion parameters using the residual background reads.

Most fragment size bias is removed thanks to our novel data structure but we still find a small amount of bias (Fig. S4). We correct for this by using a LOWESS fit with the residual background and then correct the expected background level within each region using the learned regression function.

The fitted negative binomial model represents the data well after fragment-size bias correction with a p-value distribution that is close to uniform away from zero, as expected, with a spike close to zero corresponding to data inferred to be within peaks (Fig. S5). We then use the Benjamini-Hochberg false discovery rate (FDR) correction and combined contiguously significant re-ligation sites into peaks.

### 2.3 Differential peak analysis

We take advantage of the data structure and the expected background model to develop an addition to our main software. To call differentially bound regions we first combine the peaks from all the samples to create a list of consensus peaks. Similarly to DiffBind (Stark and Brown, 2011; Ross-Innes *et al*., 2012), we then count how many reads were assigned to those regions from each sample, correct the values by removing the expected background based on the negative binomial model and fragment size and then analyse the results using DESeq2 (Love *et al*., 2014) to normalize the read counts across samples and perform differential expression analysis.

Unsupervised hierarchical clustering was done using Euclidean distance on the signal from all the peaks with rlog normalization. Motif enrichment analysis was done using differentially bound peaks between Tregs and naïve T cells and Th17 and naïve T cells. These regions were submitted to HOMER v 4.8.3 (Heinz *et al*., 2010) with the findMotifsGenome.pl command and “−size given” parameter.

### 2.4 Data pre-processing

HiChIP data from Mumbach et al. (2017) was downloaded from SRA for Naïve T cells, Th17 and Tregs (SRP# SRP112520). Reads were filtered and the adapters were removed using fastp v0.19.4. The reads were then mapped to the GRCh38 genome with HiC-Pro v2.11.0, using default settings. Replicates were merged together as described in the results section.

### 2.5 Hichipper peak calling

We called anchors using the Hichipper v 0.7.5 pipeline (Lareau and Aryee, 2018) on the HiC-Pro results with default settings making sure to include the modified background correction and restriction fragment aware padding. We called peaks with the setting EACH, SELF (for self-circle and dangling ends only) or EACH, ALL (for all reads).

### 2.6 FitHiChIP (MACS2 short range) peak calling

We used the supplied tool with FitHiChIP (Bhattacharyya *et al*., 2018) to call peaks from HiC-Pro results with default settings. This tool uses all reads from dangling ends, re-ligation and self-circle pairs and also all reads within 1kb from the valid pairs and supplies all the reads to MACS2 2.1.1 (Zhang *et al*., 2008) for peak calling.

### 2.7 Peak calling comparison

We downloaded reference H3K27ac tracks for GM12878 cells from the Encode website (accession # ENCSR000AKC), replicated peak set (accession # ENCFF367KIF).

For the Naïve T cells we used the processed peaks from the roadmap project (Sample E038) (Kundaje *et al*., 2015). We used the tool LiftOver to convert the genomic coordinates from hg19 to hg38.

All comparisons were done using bedtools v2.27.1 (Quinlan and Hall, 2010) annotate function and then analysed in python. No extension of the peaks was done. Peaks on X and Y chromosomes were excluded from the comparison. For HiChIP-peaks plots are presented as lines that result in cumulative sum of the results with the peaks sorted by p value.

### 2.8 Subsampling analysis

We created subsampled datasets from the GM12878 HiChIP data (Mumbach *et al*., 2017) by subsampling the raw reads creating datasets with 500, 250, 125 and 62.5 million reads. For the Naïve T cells dataset, we used the combined data, the 2 biological replicates with the 2 technical combined or the 4 technical replicates as individual samples.

We compared how many of the peaks called using the full dataset could be recovered from the subsampled datasets for Hichipper and HiChIP-peaks. We counted the number of peaks identified using bedtools v2.27.1 intersect in “–wa” mode and calculated the recall rate.

For loop calling we used Hichipper with default settings. We either used the default peak calling algorithm from Hichipper or we supplied the peaks called using HiChIP-peaks from the respective dataset. In the former case we used the skip-resfrag-pad setting to avoid Hichipper expanding the peaks.

To compare the loops called we first filtered the loops by FDR < 0.10 as reported by Mango (Phanstiel *et al*., 2015). We then checked if the loops called in the full dataset could be found in the subsampled datasets and calculated the recall rate. A loop was considered recalled if both ends overlapped both ends of a loop in the subsampled dataset.

## 3 Results

### 3.1 HiChIP-peaks can successfully recover reference peaks from HiChIP datasets at a higher sensitivity and specificity than previously available software

To evaluate the performance of our peak calling algorithm we chose two of the cell lines reported by Mumbach et al. (2017) for which a reference ChIP-seq track was available either from ENCODE or Roadmap project. We combined all the reads from different replicates from Naïve T cells and from GM12878 cells, respectively. Using different metrics, we show that our method is superior to previous attempts at calling peaks and allows for scoring of the peaks identified.

Specifically, our method is able to recover more peaks from the reference with significantly lower false discovery rate (Fig. 2A-B) and calling fewer peaks (Fig. S6) than Hichipper or FitHiChIP (note real false discovery rate cannot be zero because the reference ChIP-Seq does not come from the same sample as the HiChIP). In particular, we note that both Hichipper with all reads and FitHiChIP present significant false discovery rate problems with more than 70% of peaks called not observed in the reference.

**Fig. 2.**
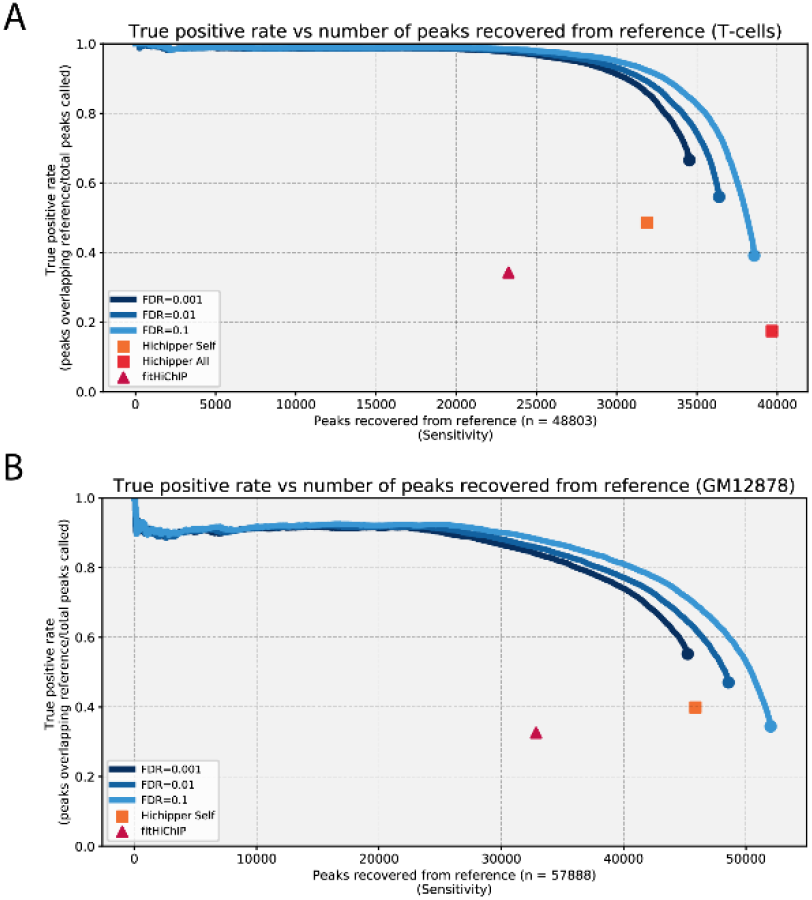
True positive rate vs number of peaks recalled from reference in (A) T-cells and (B) GM12878 cells. For our software we sort peaks in ascending p-value order and show the true positive rate as the number of peaks recovered increases. We provide three different FDR settings as the FDR setting changes the size of the peaks themselves and the lines do not overlap perfectly. We show results for Hichipper and fitHiChIP (default settings) for comparison. Hichipper in (ALL) mode fails to run with the GM12878 dataset.

Moreover, we note that with Hichipper it is not possible to change the sensitivity: changing the q-value setting doesn’t produce any difference in number of peaks called or genome covered.

### 3.2 HiChIP-peaks is more stable than Hichipper when read depth is reduced

Using the best settings for Hichipper (SELF reads) we compared the stability of the results when the number of reads in the dataset is reduced. We analysed the individual technical replicates of the Naïve T cells that contain about 100 million reads per sample. We show that our method is consistently able to maintain accuracy and sensitivity, while Hichipper suffers greatly when the number of reads is less than optimal (Fig. 3). We then tested how many of the peaks from a larger dataset could be recalled from a smaller dataset. We used progressively subsampled datasets for the GM12878 dataset and we tested technical replicates and biological replicates from the naïve T cell dataset. HiChIP-peaks has a much higher recall rate in both of these cell types when the read count is reduced compared to Hichipper (Fig. 4 A, C).

**Fig. 3.**
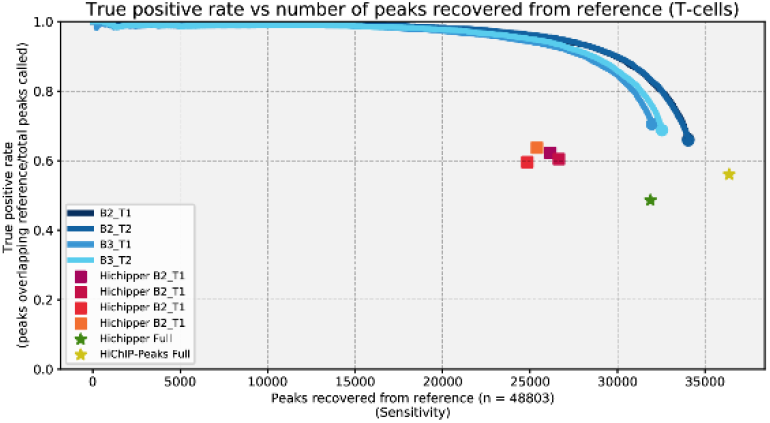
Effect of reduced read depth on peak calling performance. True positive rate vs number of peaks recalled from reference (Naïve T cells dataset). We show that our software maintains high consistency while Hichipper’s sensitivity goes down rapidly when read count goes down. Our software is set at a FDR of 0.01.

**Fig. 4.**
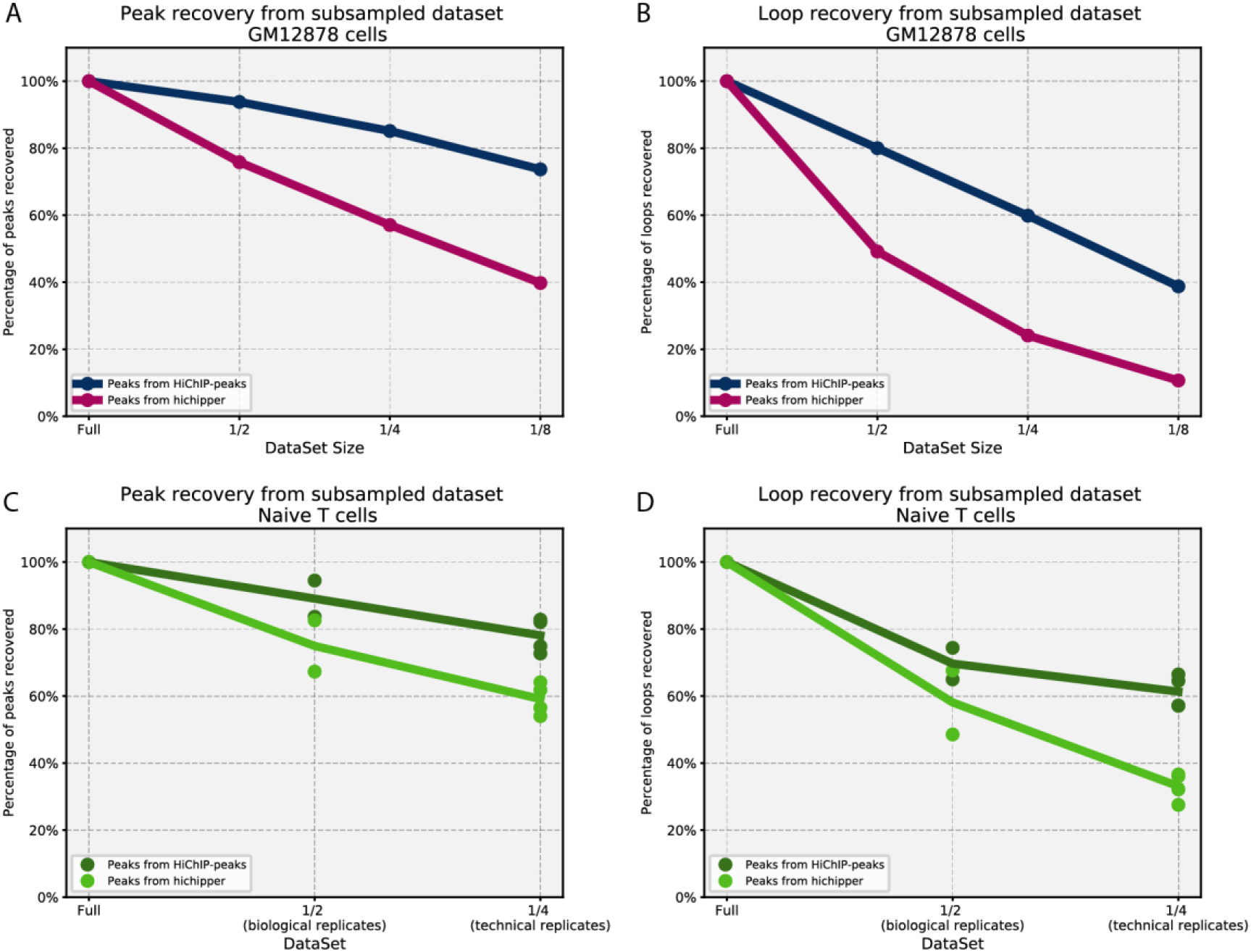
Effect of subsampled datasets on peak and loop calling. A) Recall rate of peaks from full dataset using subsampled datasets from GM12878 dataset. B) Recall rate of loops called (chr22) from full dataset using GM12878 datasets. C) Same as A but with Naïve T cells dataset. We merged the technical replicates for each biological replicates for ½ and we used each technical replicate individually for ¼. D) same as B but with Naïve T cells datasets.

### 3.3 Peaks can be supplied to Hichipper for analysis and provide better consistency when lowering dataset depth

We decided to test whether our improved peak calling would affect loop calling results from Hichipper. Because the biggest differences were found when lowering the number of reads we decided to test Hichipper’s stability using progressively subsampled datasets from the GM12878 dataset and by combining fewer technical replicates from the naïve T cells datasets. We note that our algorithm is able to produce a higher recall rate compared to Hichipper and we see that using peaks called with our algorithm yields significantly better stability in recall of loops for both cell types as the datasets size is reduced (Fig. 4 B, D). This shows that stability and accuracy of the peaks called significantly impacts the loop calling results and our algorithm can greatly improve the stability of the results, especially when number of reads available is limited.

### 3.4 Novel data representation allows accurate differential peak calling

Using the novel data representation, we provide an interface to analyse differentially bound regions in HiChIP datasets, fully exploiting the information contained in them.

We carried out a proof-of-concept study by analysing the four technical replicates of the naïve T cells individually. Our results show that the sensitivity and reproducibility of our software is sufficiently good that we can easily differentiate between technical and biological replicates of the same cell type (Fig. 5 A). We find almost 3000 peaks (more than 10% of all peaks) differentially bound (FDR<0.10, log2FoldChange>0.5) between biological replicates of the same cell type further affirming the importance of peak calling on individual HiChIP datasets instead of using combined or external ChIP-seq datasets.

**Fig. 5.**
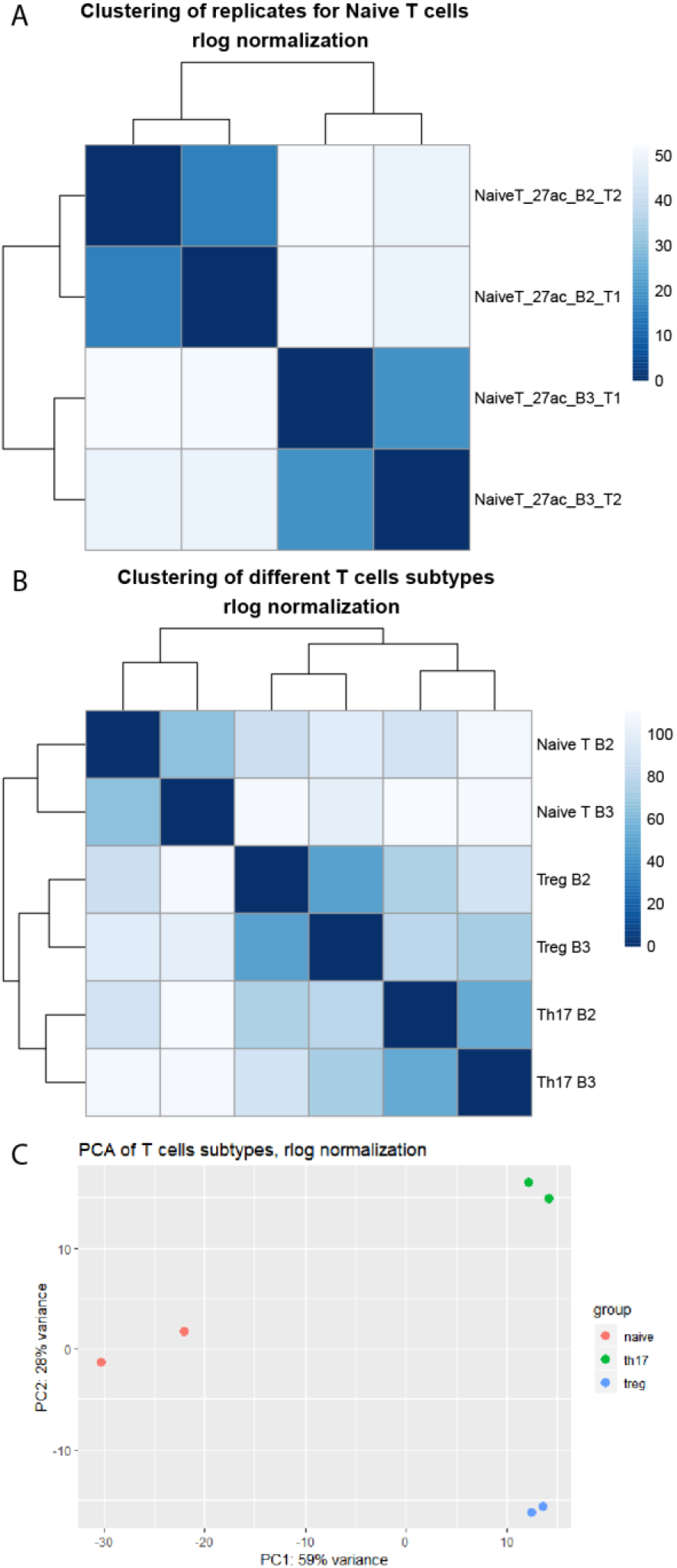
Differential peak analysis. A) Unsupervised hierarchical clustering using scores from the 4 technical replicates of Naïve T cells. B) Unsupervised hierarchical clustering of the 3 different T-cell related cell lines. C) PCA of the 3 different cell lines.

We then analysed data from the two other T cell types, Th17 and Tregs. We merged the technical replicates into biological replicates. Although the read depth is very different between the different samples (37m to 60m reads used in the peak calling) our software performs remarkably consistently. Biological differences vastly outweigh technical differences and samples cluster by cell type (Fig. 5 B, C). We identify thousands of peaks that are significantly differentially bound between the different cell types. As expected the differences between Th17 and Tregs are smaller than between Th17 and Naïve T cells. To test whether the differentially bound peaks have biological significance we ran motif enrichment analysis with HOMER on the peaks from Th17 vs Naïve T cells and Tregs vs Naïve T cells contrasts. The results clearly indicate enrichment in binding sites for transcription factors involved in the interferon pathway, ETS-RUNX and others (Table S2), consistent with models of T cell activation (Christie and Zhu, 2014) and confirming the accuracy of our peak calling method.

## 4 Discussion

HiChIP is quickly gaining importance, especially in studies involving primary cells of various tissue types thanks to the lower input and sequencing requirements. Previously ChIP-seq tracks were used to identify peak regions as the quality of peak calling from HiChIP was deemed insufficient. This either added a significant cost and sample requirement to the experiment design or often researchers relied on data not generated from the same sample.

We show that our software can reliably and efficiently identify enriched regions using only HiChIP datasets, even when read depth is relatively low, and that using HiChIP-peaks significantly improves the reliability of Hichipper’s loop calling. This shows also how good peak calling is of fundamental importance for Hichipper’s functionality. Our results also demonstrate that accurate peak calling from each sample is important because each biological replicate can have different peaks, which can affect the identified loops, especially when studying more transient and regulated regions.

As the popularity of chromatin conformation methods increase, commercial kits, such as Arima Hi-C, are starting to be developed. This kit, with modifications in the protocol, can be used to generate HiChIP libraries. The kit is highly efficient thanks to its dual restriction enzyme protocol, but this results in the absence of reported dangling ends and self-circles (Fig. S2). This impacts the performance of Hichipper using the SELF setting, which, according to our analysis, is the best of the currently available methods (Fig 2). Therefore, our method, HiChIP-Peaks, has the potential to be the only method of choice when using commercially available kits such as Arima Hi-C to generate HiChIP libraries.

Our results show that our alternative data structure for representing Hi-C reads limits biases due to how reads are generated in this protocol and maximises resolution within the constraints of the technology. This data structure can also be used for other kinds of analysis with simple generalizations.

## Acknowledgements

The authors would like to acknowledge the assistance given by IT Services and the use of the Computational Shared Facility at The University of Manchester.

## Supplementary tables and figures

**Table S1.** Number of reads that contain uncut restriction sites within HiChIP datasets. Approximately 15 million out of the 98 million reads from the dataset contain an uncut MboI restriction site motif. In dark blue are the 6bp motifs that could have been generated by the re-ligation event. All other motifs are evidence of uncut restriction sites. Dataset from Naïve T cells, Biological replicate 2, technical replicate 1, read 1 (Fwd).

| Accession number | Reads |  |
| --- | --- | --- |
| SRR5831497 (Fwd) | 98201048 | Approximately 15 million uncut restriction sites on R1 |

| Motif | counts | Motif | counts |
| --- | --- | --- | --- |
| AGATCA | 1139756 | TGATCA | 840523 |
| AGATCC | 738643 | TGATCC | 1062812 |
| AGATCT | 875481 | TGATCT | 1167673 |
| AGATCG | 12627791 | TGATCG | 11874109 |
| CGATCA | 11220492 | GGATCA | 1154531 |
| CGATCC | 10110047 | GGATCC | 660955 |
| CGATCT | 11952588 | GGATCT | 741551 |
| CGATCG | 3949528 | GGATCG | 11027058 |

**Table S2.** Motifs identified as significantly enriched from differentially bound peaks. Results from HOMER using differentially bound peaks from the specified contrasts.

Tregs vs Naïve T cells
| Motif Name | P-value | Log P-value | q-value (Benjamini) |
| --- | --- | --- | --- |
| ISRE(IRF)/ThioMac-LPS-Expression(GSE23622) | 1.00E-21 | -5.00E+01 | 0 |
| IRF2(IRF)/Erythroblas-IRF2-ChIP-Seq(GSE36985) | 1.00E-17 | -4.06E+01 | 0 |
| IRF1(IRF)/PBMC-IRF1-ChIP-Seq(GSE43036) | 1.00E-10 | -2.33E+01 | 0 |
| Ets1-distal(ETS)/CD4+-PolII-ChIP-Seq(Barski_et_al.) | 1.00E-08 | -1.85E+01 | 0 |
| Bach1(bZIP)/K562-Bach1-ChIP-Seq(GSE31477) | 1.00E-06 | -1.45E+01 | 0 |
| NF-E2(bZIP)/K562-NFE2-ChIP-Seq(GSE31477) | 1.00E-06 | -1.39E+01 | 0.0001 |
| ETS:RUNX(ETS,Runt)/Jurkat-RUNX1-ChIP-Seq(GSE17954) | 1.00E-05 | -1.34E+01 | 0.0001 |
| Nrf2(bZIP)/Lymphoblast-Nrf2-ChIP-Seq(GSE37589) | 1.00E-04 | -1.09E+01 | 0.0008 |
| Jun-AP1(bZIP)/K562-cJun-ChIP-Seq(GSE31477) | 1.00E-03 | -8.39E+00 | 0.0088 |
| E2F(E2F)/Hela-CellCycle-Expression | 1.00E-03 | -7.29E+00 | 0.024 |
| RFX(HTH)/K562-RFX3-ChIP-Seq(SRA012198) | 1.00E-03 | -6.92E+00 | 0.0318 |
| NFkB-p65-Rel(RHD)/ThioMac-LPS-Expression(GSE23622) | 1.00E-02 | -6.75E+00 | 0.035 |

Th17 vs Naïve T cells
| Motif Name | P-value | Log P-value | q-value<br>(Benjamini) |
| --- | --- | --- | --- |
| ISRE(IRF)/ThioMac-LPS-Expression(GSE23622) | 1.00E-11 | -2.66E+01 | 0 |
| IRF2(IRF)/Erythroblas-IRF2-ChIP-Seq(GSE36985) | 1.00E-09 | -2.29E+01 | 0 |
| ETS:RUNX(ETS,Runt)/Jurkat-RUNX1-ChIP-Seq(GSE17954) | 1.00E-08 | -1.85E+01 | 0 |
| Ets1-distal(ETS)/CD4+-PolII-ChIP-Seq(Barski_et_al.) | 1.00E-08 | -1.84E+01 | 0 |
| NFkB-p65-Rel(RHD)/ThioMac-LPS-Expression(GSE23622) | 1.00E-04 | -1.14E+01 | 0.0008 |
| RFX(HTH)/K562-RFX3-ChIP-Seq(SRA012198) | 1.00E-04 | -1.01E+01 | 0.0022 |
| Bach1(bZIP)/K562-Bach1-ChIP-Seq(GSE31477) | 1.00E-04 | -9.50E+00 | 0.0036 |
| IRF1(IRF)/PBMC-IRF1-ChIP-Seq(GSE43036) | 1.00E-03 | -9.17E+00 | 0.0045 |
| Nrf2(bZIP)/Lymphoblast-Nrf2-ChIP-Seq(GSE37589) | 1.00E-03 | -7.84E+00 | 0.0152 |
| Rfx2(HTH)/LoVo-RFX2-ChIP-Seq(GSE49402) | 1.00E-03 | -7.45E+00 | 0.0205 |
| NF-E2(bZIP)/K562-NFE2-ChIP-Seq(GSE31477) | 1.00E-03 | -7.12E+00 | 0.026 |
| T1ISRE(IRF)/ThioMac-Ifnb-Expression | 1.00E-02 | -6.63E+00 | 0.0394 |

**Figure S1.**
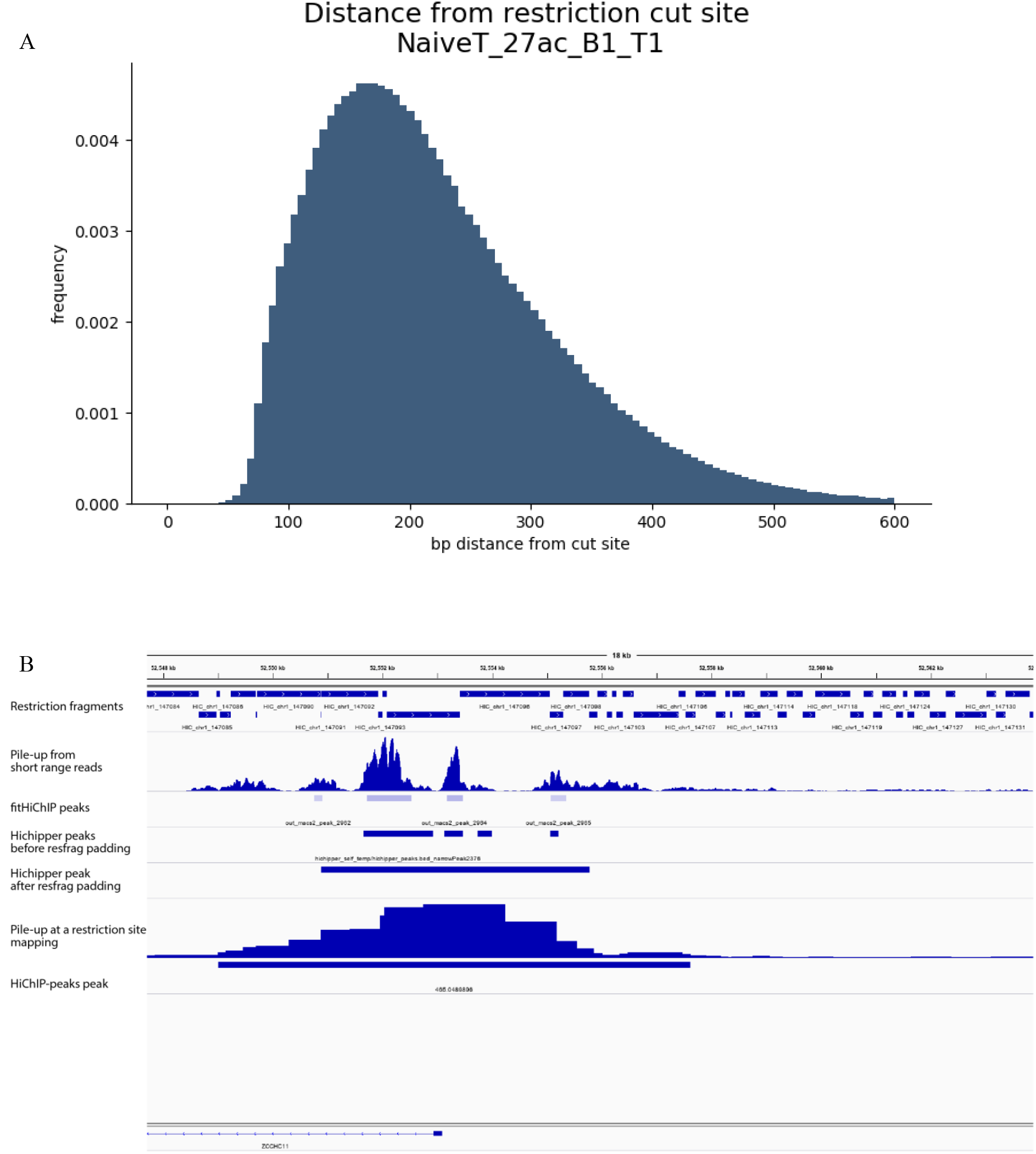
Reads are located around the restriction site, as expected from a Hi-C library. A) Distribution of reads’ distance from closest restriction site. Dataset from Naïve T cells, Biological replicate 1, technical replicate 1. B) Pileup of short-range reads from HiChIP dataset Naïve T cells combined (using FitHiChIP’s utility). The signal is more intense around the restriction sites, creating sparsity elsewhere which can bias other peak calling methods resulting in many small peaks. Hichipper attempts to compensate this by extending the peaks to the nearest fragment. HiChIP-peaks is based on the re-ligation site and doing so ignores the sparsity by design and maximises usable information.

**Figure S2.**
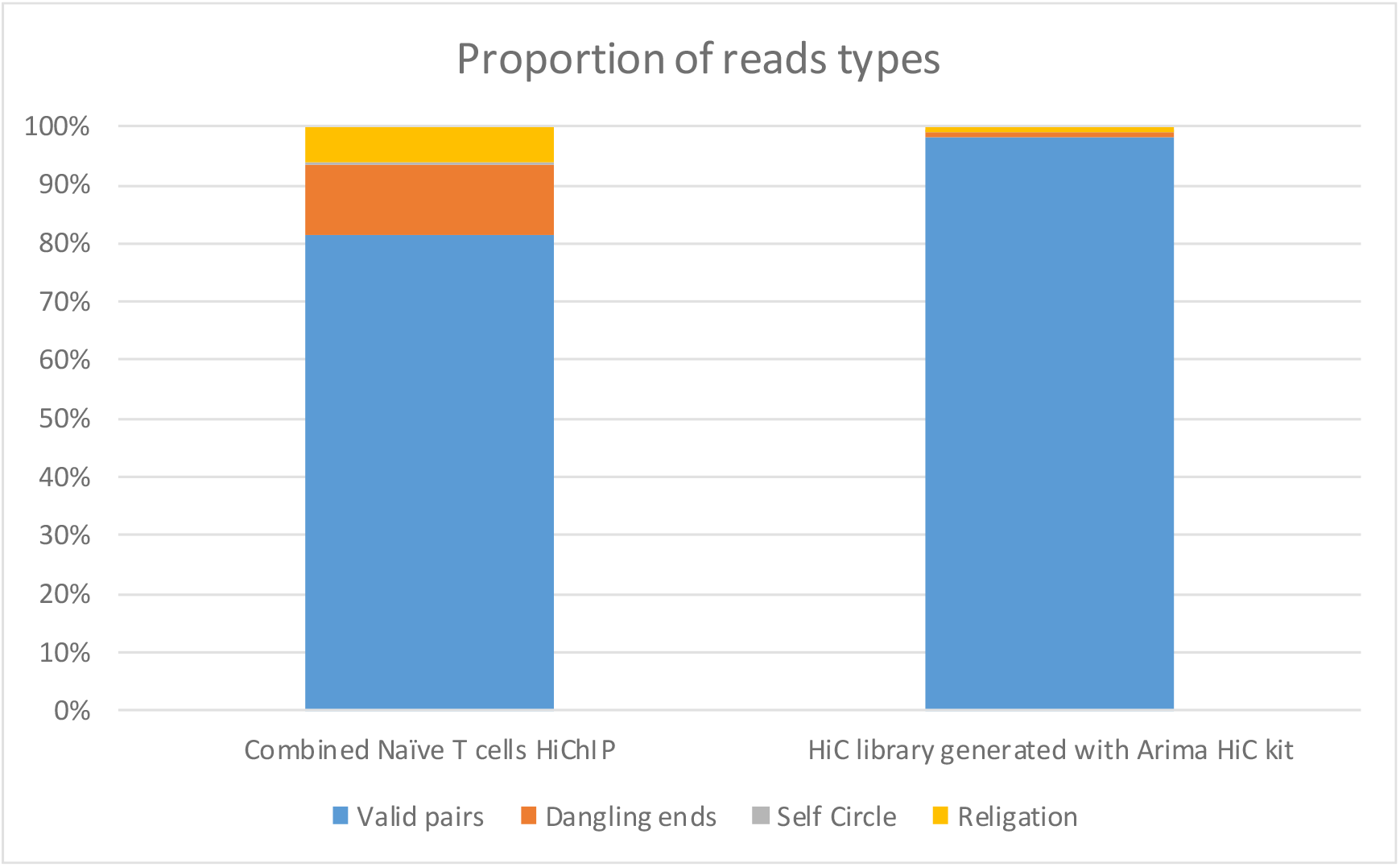
Proportion of read pairs identified as dangling ends, self-circle and re-ligation from the Naïve T cells dataset and a Hi-C library generated with the Arima-Hi-C kit (unpublished). Libraries generated with the Arima-Hi-C kit have almost no dangling ends and self-circle reads that can be used by Hichipper in SELF mode.

**Figure S3.**
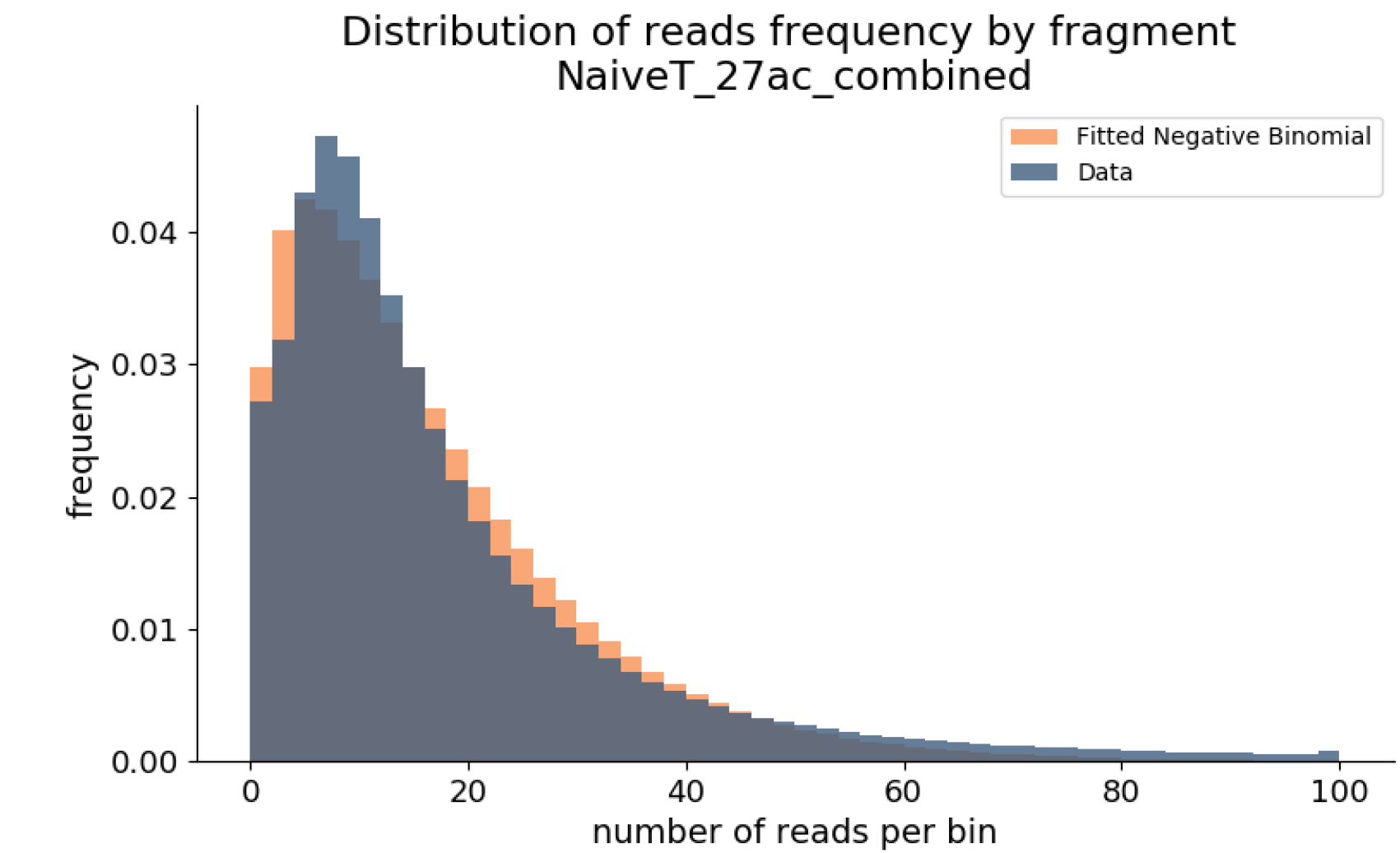
Read count distribution per restriction site as used for HiChIP-Peaks. The noise signal from the HiChIP datasets resembles a negative binomial distribution. Data from the combined Naïve T cells dataset.

**Figure S4.**
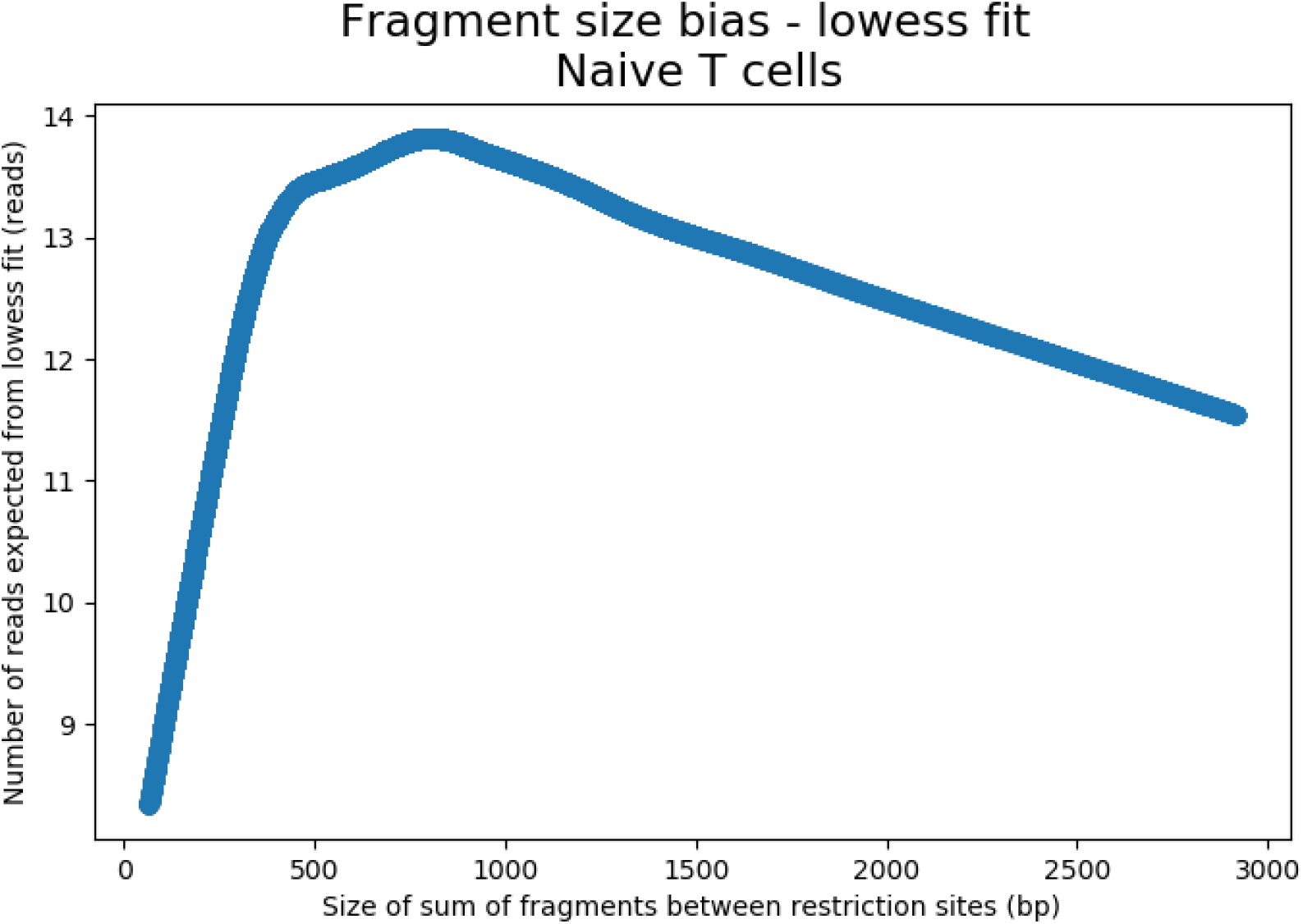
Fragment size bias fitted using a LOWESS fit. Fragment size is the sum of the fragments within the tested re-ligation sites. We use this fit to correct the expected background by fragment size in the negative binomial test. Data from the combined Naïve T cells dataset.

**Figure S5.**
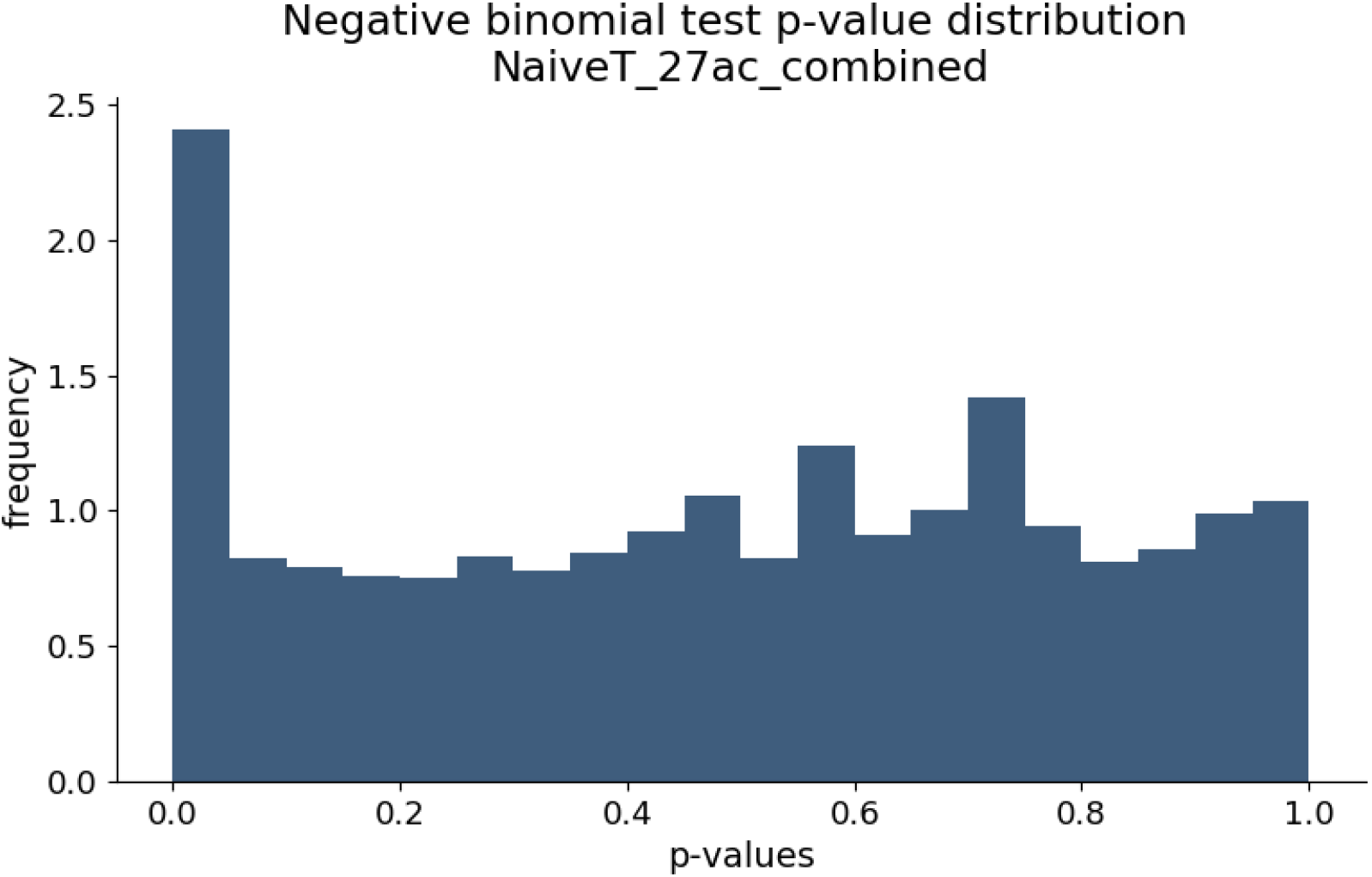
P-value distribution from negative binomial test. The null distribution is uniform, showing an appropriate fit of the background model. Data from the combined Naïve T cells dataset.

**Figure S6.**
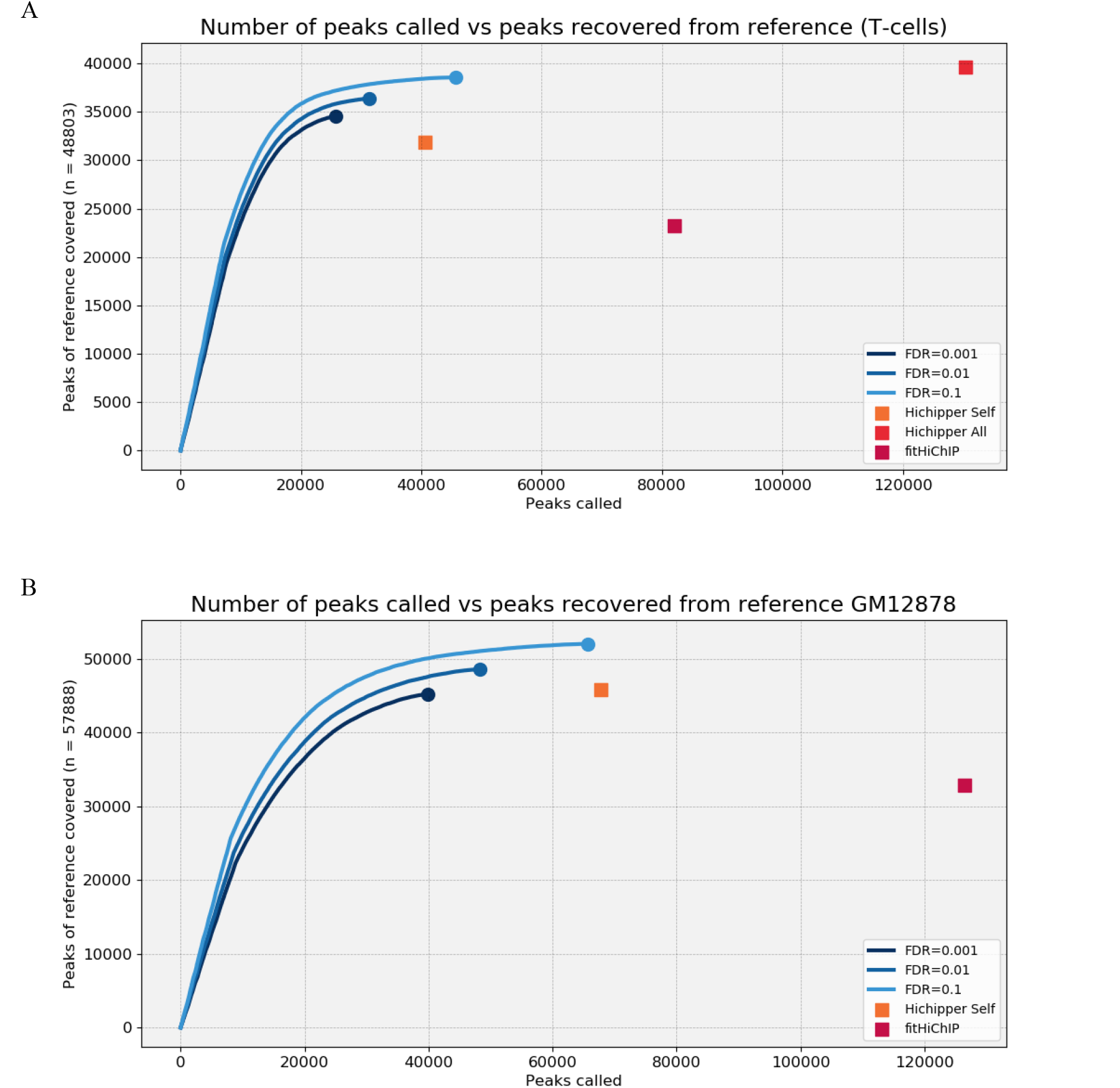
Peaks recalled from reference vs peaks called from Naïve T cells (A) and GM12878 (B). Our algorithm can identify more peaks from the reference while calling fewer peaks.

